# Integrating redox dynamics and gene networks reveals ROS-mediated control of Arabidopsis seed germination under water stress

**DOI:** 10.64898/2026.11.16.766966

**Authors:** Xiujie Mu, Adrien Vidal, Stephan Wagner, Andreas Meyer, Fredy Barneche, Christophe Bailly

## Abstract

Seed germination is a critical step of plant development that is highly sensitive to water availability. During this delicate transition, water stress responses are tightly regulated by multilevel molecular networks that integrate environmental cues into developmental adaptations. Reactive Oxygen Species (ROS) play a positive signaling role in seed germination, yet their influence on germination under water stress remains poorly understood. Here, we investigated the involvement of ROS in *Arabidopsis thaliana* seed germination upon reproducibly reducing water potential using polyethylene glycol (PEG). The data show that delayed germination under water stress is not simply a consequence of reduced water uptake. Instead, water stress triggers coordinated reprogramming events characterized by altered H_2_O_2_ homeostasis, particularly in peroxisomes, which functionally mediates a rebalancing of ABA and GA metabolism toward developmental slowdown. A weighted gene co-expression network analysis of water-stress transcriptomic dynamics identified a gene hub associated with H_2_O_2_ production during germination, shedding light on multiple oxygen-and redox-related processes that are strongly linked to transcriptional and translational attenuation. Collectively, this study unveils a role for regulating peroxisomal H_2_O_2_ homeostasis in *Arabidopsis* seed water stress responses, offering new perspectives for improving crop resilience to environmental constraints.

## INTRODUCTION

Seed germination is a critical stage of plant development that is highly sensitive to water availability. Water deficit significantly reduces germination speed, delays radicle protrusion, and decreases seedling establishment (Vallejo et al., 2010; Liu et al., 2019; Wang et al., 2022; Kolesnikov et al., 2023). The importance of germination for crop productivity is illustrated by numerous studies showing that water deficit during or immediately after sowing reduces emergence rates, delays establishment, and ultimately decreases yield in major crops, including wheat, maize, rice, soybean, sunflower, and rapeseed (Farooq et al., 2009; Liu et al., 2010; Hasanuzzaman et al., 2013; Bailly and Gomez Roldan, 2023). Because seedling establishment represents a major bottleneck in crop production, improving germination performance under water stress is increasingly recognized as a priority for crop adaptation to climate change (Bailly and Gomez Roldan, 2023). Although the inhibitory effects of water stress on germination have been widely documented, the molecular mechanisms through which germinating seeds perceive water limitation and translate this environmental cue into coordinated changes in hormonal and developmental responses remain poorly understood.

Reactive Oxygen Species (ROS) are attractive candidates for constituting the initial sensing layer during germination under water-limited conditions. According to the “oxidative window” concept, successful germination requires ROS accumulation within a permissive range that is sufficient to activate germination-related processes without causing oxidative damage (Bailly et al., 2008). Hence, rather than acting solely as toxic by-products of aerobic metabolism, ROS are broad integrators of environmental signals during germination (Bailly, 2008, 2019, 2022, Jurdak et al., 2020). ROS signaling output depends on their concentration, subcellular localization, and temporal dynamics. In seeds of sunflower, water stress induces significant changes in ROS accumulation and antioxidant enzyme activities during germination (Saux et al., 2020), while similar responses have been reported in seedlings exposed to drought conditions (Vallejo et al., 2010; Liu et al., 2019; Wang et al., 2022; Kolesnikov et al., 2023). Beyond their overall abundance, the cellular origin and subcellular distribution of ROS are increasingly recognized as critical determinants of their signaling functions. During germination, ROS are notably generated in mitochondria, peroxisomes, the apoplast, and plasma membrane-associated NADPH oxidases, each contributing distinct redox signals (Bailly, 2019; Jurdak et al., 2022). Among these compartments, peroxisomes are of particular interest because they are highly active during germination and play central roles in reserve mobilization through fatty acid β-oxidation and the glyoxylate cycle (Graham, 2008). These metabolic pathways generate H_2_O_2_ as a by-product, positioning peroxisomes at the interface between metabolism and redox signaling (Graham, 2008; Pracharoenwattana et al., 2010). Peroxisomes may therefore act not only as metabolic organelles but also as important regulators of germination through ROS production (Jurdak et al., 2022). However, despite increasing recognition of the importance of ROS compartmentation, little is known about how environmental stresses influence ROS dynamics within specific organelles during germination. In particular, whether water stress modifies peroxisome-derived ROS signaling and how such changes contribute to the regulation of germination progression remain largely unexplored.

ROS are likely to influence germination by modulating the metabolism and signaling pathways of the phytohormones abscisic acid (ABA) and gibberellic acid (GA) (Bailly, 2022), which directly contribute to germination-associated water stress responses. Under water deficit, ABA levels and signaling generally increase, reinforcing dormancy-related processes and inhibiting germination, whereas GA biosynthesis and responsiveness are reduced, limiting embryo growth and radicle protrusion (Llanes et al., 2016; Liu et al., 2019; Hassan et al., 2021). In *A. thaliana*, rice, sunflower, and wheat, low water potential conditions induce genes associated with ABA signaling, osmotic stress responses, antioxidant systems, and reserve mobilization (Liu et al., 2019; Saux et al., 2020; Kolesnikov et al., 2023). Increasing evidence indicates that ROS are closely integrated into this hormonal network. Hydrogen peroxide (H_2_O_2_) addition promotes germination by stimulating GA biosynthesis and ABA catabolism while repressing ABA biosynthesis and GA degradation (Liu et al., 2010; Cheng et al., 2022). Conversely, ABA suppresses ROS accumulation in several species, including *A. thaliana*, notably through the activation of the ABA signaling transcription factor ABA-INSENSITIVE_5 (ABI5), which induces ROS scavenging via CATALASE1 (Ye et al., 2012; Lariguet et al., 2013; Bi et al., 2017). Together, these observations indicate that ROS are important intermediates in linking water stress perception to the transcriptional regulation of ABA-GA signaling and germination progression.

In vegetative tissues, drought similarly induces genes involved in ABA signaling, ROS detoxification, osmoprotection, and stress defense, while repressing pathways associated with cell division, protein synthesis, and growth (Shinozaki and Yamaguchi-Shinozaki, 2006; Skirycz and Inzé, 2010; Claeys and Inzé, 2013; Nakashima et al., 2014). At the molecular level, these responses involve large-scale activation of ABA-responsive transcription factors, including members of the AREB/ABF, NAC, DREB, and WRKY families, together with extensive reorganization of primary metabolism and redox homeostasis (Shinozaki and Yamaguchi-Shinozaki, 2006; Nakashima et al., 2014). Notably, drought-induced growth inhibition is frequently associated with repression of ribosome biogenesis, translation, DNA replication, and chromatin-associated functions, suggesting that plants actively redirect resources from growth toward stress adaptation (Skirycz and Inzé, 2010; Claeys and Inzé, 2013). Compared with vegetative tissues, much less is known about transcriptomic responses to water stress during seed germination, and a comprehensive understanding of how water stress triggers a coordinated, temporally regulated reprogramming linking redox signaling, hormonal responses, and developmental competence during germination remains elusive.

We hypothesized that water stress modifies ROS homeostasis and associated signaling pathways, thereby altering the regulatory networks controlling germination progression. Combining germination phenotyping, transcriptomic analyses, ROS biochemical measurements and *in vivo* monitoring of subcellular redox dynamics, we deciphered how water stress affects ROS homeostasis and associated signaling pathways across successive stages of germination. Our study identifies the temporal sequence of molecular events that link water stress perception to delayed germination and reveals a central role for peroxisome-associated ROS signaling in these regulatory mechanisms.

## RESULTS

### Seed germination under water stress triggers transcriptome reprogramming

*A. thaliana* wild-type seeds were germinated on PEG solutions with water potentials ranging from -0.2 MPa to -1.2 MPa. While stratified seeds fully germinated within 3 days in water, increasing water potential gradually delayed germination (Figure 1A). For the highest level tested (-1.2 MPa), the final germination rate was severely reduced (Figure 1A). For the remainder of this study, the -0.9 MPa PEG condition was selected because it delayed germination without affecting the final germination percentage (Figure 1A). Based on the germination curves (Figure 1A), three sampling time points corresponding to successive stages of the progression toward germination completion were defined for each condition (Figure 1B). These included an early stage (ET, 3 h after imbibition), a middle stage (MT, corresponding to half of the time required to reach the late stage), and a late stage (LT), defined as one day before the onset of radicle protrusion. Although the time required to reach these stages differed with each water potential tested, the selected sampling points represented equivalent physiological stages of germination progression under both conditions (Figure 1B). The water stress imposed by the -0.9 MPa PEG solution can be considered moderate, as seed water content decreased only slightly, from approximately 0.83 to 0.80 g H_2_O/g fresh weight, during imbibition (Figure 1C), and radicle emergence was not prevented.

**Figure 1.**
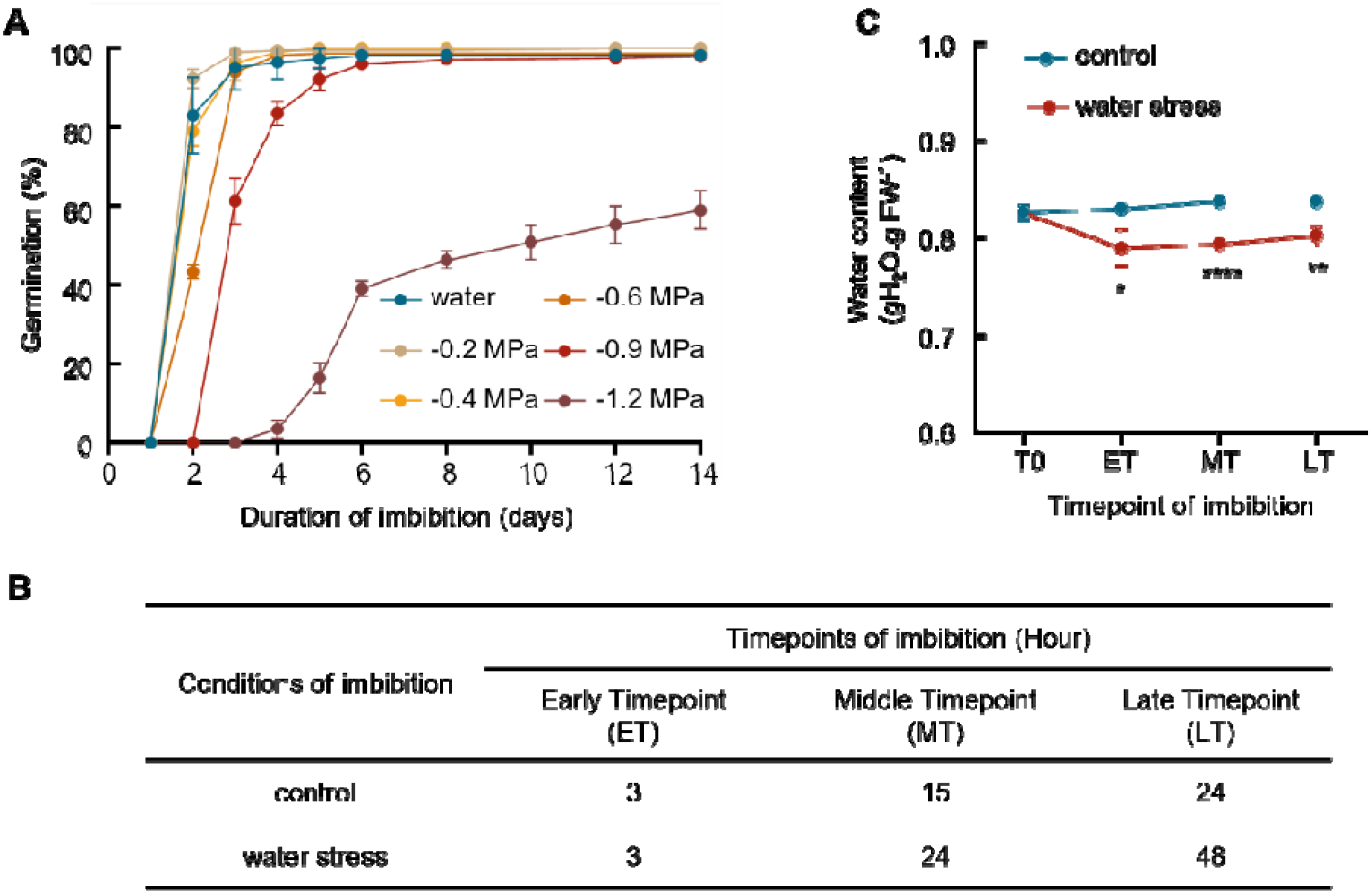
Water stress delays seed germination. (**A**) Germination of stratified seeds in water or a range of PEG solutions at 15°C in the dark (**B**) The 3 time points used for analyses, defined according to the germination kinetics on water and on a -0.9 MPa PEG solution (**C**) Seed water content at initial time (after stratification, T0) and at the 3 timepoints defined in Figure 1B during imbibition at 15°C under the dark. Data presented are means of 3 replicates ± SD. Student’s t-test was used to determine the statistical significance between the two groups at each time point. An asterisk below the bar chart indicates a significant difference between the two groups (*, P < 0.05; **, P < 0.01; ***, P < 0.001).

To investigate transcriptomic responses to water stress during germination, stratified seeds were imbibed in water or in a PEG solution at -0.9 MPa and samples were used for RNA-seq analyses at the 3 time points defined above. Water stress triggered 252, 9844, and 12,653 differentially expressed genes (DEGs) at ET, MT, and LT, respectively (Figure 2A, Table S2-4). At ET, upregulated genes were enriched in secondary metabolism, sulfur compound biosynthesis, glucosinolate metabolism, and stress-related pathways, and included well-characterized stress-responsive genes (Table S1), such as *HSP17.6B*, a member of the heat shock protein (HSP) family (Chen et al., 2024), *LTP3*, encoding a non-specific lipid transfer protein (Gao et al., 2016), and *PUB19*, encoding a U-box E3 ubiquitin ligase involved in ABA and stress signaling (Liu et al., 2011). Concomitantly, several members of the xyloglucan endotransglucosylase/hydrolase family, including *XTH16*, *XTH19*, *XTH22*, and *XTH27* (Rose et al., 2002), were significantly downregulated under PEG treatment (Table S1). In addition, the glycosyltransferase gene *F8H* (Lee et al., 2009) and the growth-promoting gene *EXORDIUM* (*EXO*; Schröder et al., 2009) were repressed (Table S1). At MT, up-regulated genes were significantly enriched in GO categories related to oxygen deprivation and redox metabolism (Figure 2B), including *GPX8* (Glutathione peroxidase 8; Gaber, 2013); *GSTF11* (Glutathione S-transferase F11; Horváth et al., 2023); *GSTU14* and *GSTU2* (Glutathione S-transferase U14/2; Horváth et al., 2020), *SRO5* (Similar to RCD-ONE 5; (Babajani et al., 2009), as well as hypoxia-responsive genes, including *AHB1* (*Arabidopsis* nonsymbiotic hemoglobin 1; Perazzolli et al., 2004), *HRU1* (Hypoxia responsive universal stress protein 1; Gonzali et al., 2015), and *ADH1* (Alcohol dehydrogenase 1; Peng et al., 2001) together with members of the small heat shock protein family, were strongly induced (Table S2). Down-regulated genes were mainly associated with processes linked to RNA metabolism and seed developmental programs (Figure 2B). At LT, the transcriptional response shifted toward the regulation of signaling and enzymatic activities with up-regulated genes predominantly enriched in GO terms associated with protein phosphorylation and kinase regulation, including regulation of cyclin-dependent protein serine/threonine kinase activity, and more generally the regulation of catalytic activity (Figure 2D). The “electron transport chain” GO terms remained over-represented at this stage, suggesting a persistent adjustment of respiratory and redox-related processes under water stress. Conversely, down-regulated genes at LT were enriched in categories related to mRNA and non-coding RNA metabolism and developmental regulation (Figure 2D). Lastly, key genes involved in peroxisomal ROS production were downregulated, including *ACX1* and *ACX2*, which encode acyl-CoA oxidases catalyzing the H_2_O_2_-generating step of β-oxidation (Eastmond et al., 2000; del Río et al., 2006). Expression of the peroxin 5 (*PEX5*) gene, which encodes a major receptor required for peroxisomal protein import (Skowyra and Rapoport, 2022), was also reduced (Table S3), suggesting a broader attenuation of peroxisomal metabolic activity.

**Figure 2.**
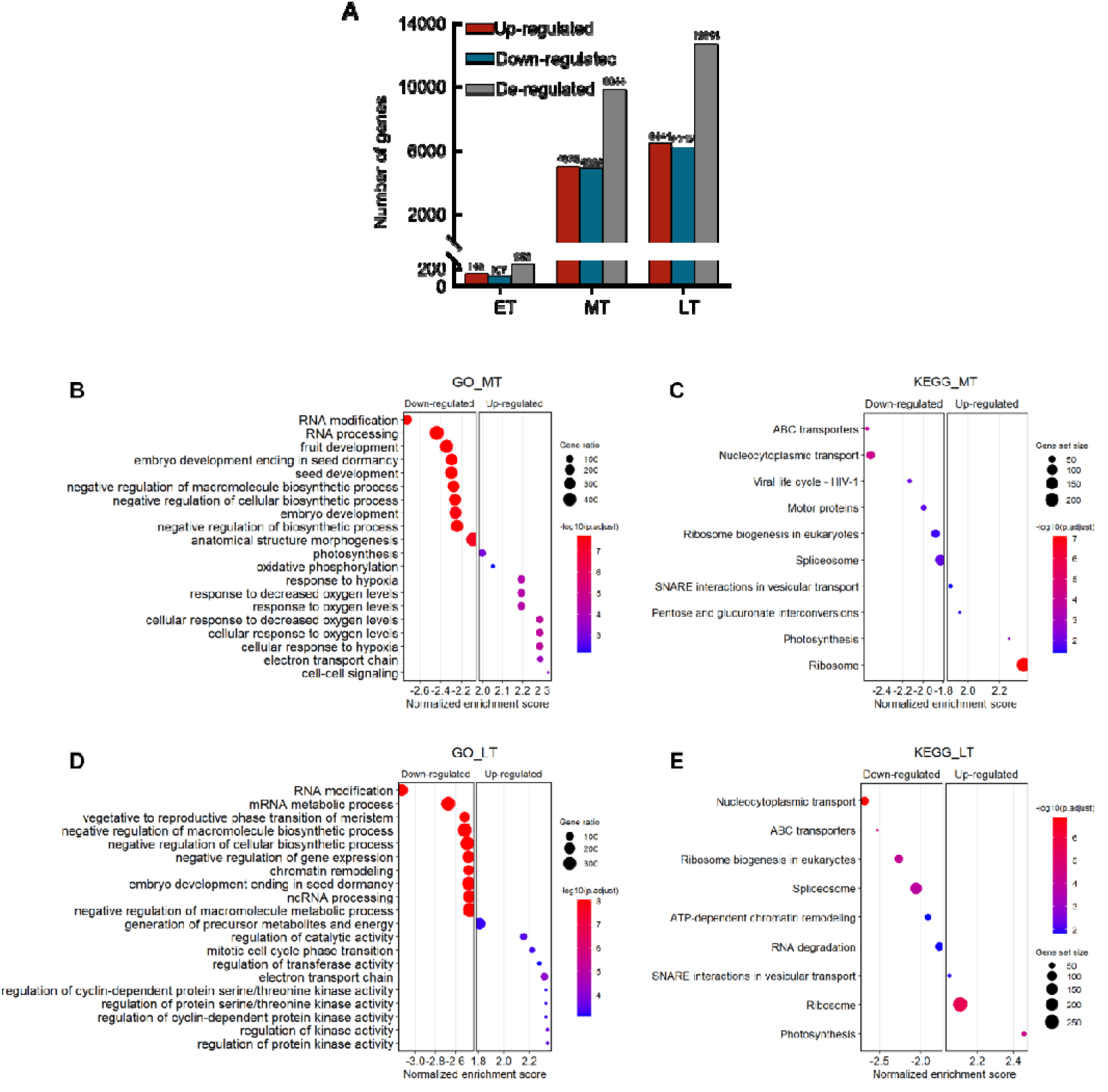
Water stress during germination induces specific seed transcriptomic responses. (**A**) Number of differentially expressed genes (DEGs), including up- and down-regulated genes (FDR < 0.01), at the 3 germination time points at 15°C under water stress conditions (ET, early timepoint; MT, mid timepoint; LT, late timepoint). (**B**-**E**) Gene set enrichment analysis (GSEA) of DEGs. The scatter plots show significantly enriched Gene Ontology (GO) terms and KEGG pathways under water stress at MT (**B**, **C**) and LT (**D**, **E**). Each dot represents a biological process or pathway, and its regulation status (up- or down-regulated) is indicated by the normalized enrichment score. Dot size represents the proportion of regulated genes relative to the total number of genes in the corresponding pathway, while color indicates the adjusted p-value.

KEGG pathway enrichment analyses further showed that, at MT, up-regulated genes are strongly enriched for pathways related to the translational machinery and RNA processing, notably ribosome and spliceosome terms, as well as photosynthesis and pentose and glucuronate interconversions (Figure 2C). By contrast, down-regulated genes were enriched in pathways associated with intracellular trafficking and transport. At LT, functions related to ribosome activity, photosynthesis, RNA degradation, and “spliceosome” pathways were still significantly upregulated, whereas down-regulated pathways included nucleocytoplasmic transport, ABC transporters, ribosome biogenesis, and ATP-dependent chromatin remodeling (Figure 2E).

Altogether, these analyses indicate that water stress induces a major transcriptional reprogramming during *Arabidopsis* seed germination, characterized by activation of oxygen-and redox-responsive pathways at MT, followed by the induction of kinase-related signaling pathways at LT, while maintaining strong modulation of ribosome-associated functions throughout germination.

### Mitochondrial electron transport chain activity is altered by water stress during seed germination

Considering that water stress specifically induces gene sets associated with mitochondrial electron transport chain (mETC), which plays a critical role in energy production during germination (Nietzel et al., 2020), we assessed seed respiratory activity during germination. As estimated by changes in oxygen uptake within a 10 min interval before and after the selected time point, respiration rate progressively decreased with increasing PEG concentrations, with the strongest inhibition observed 24 h after imbibition (Figure 3A). Consistent with these results, respiration rates were significantly reduced at the late stage of germination under -0.9 MPa (Figure 3B).

**Figure 3.**
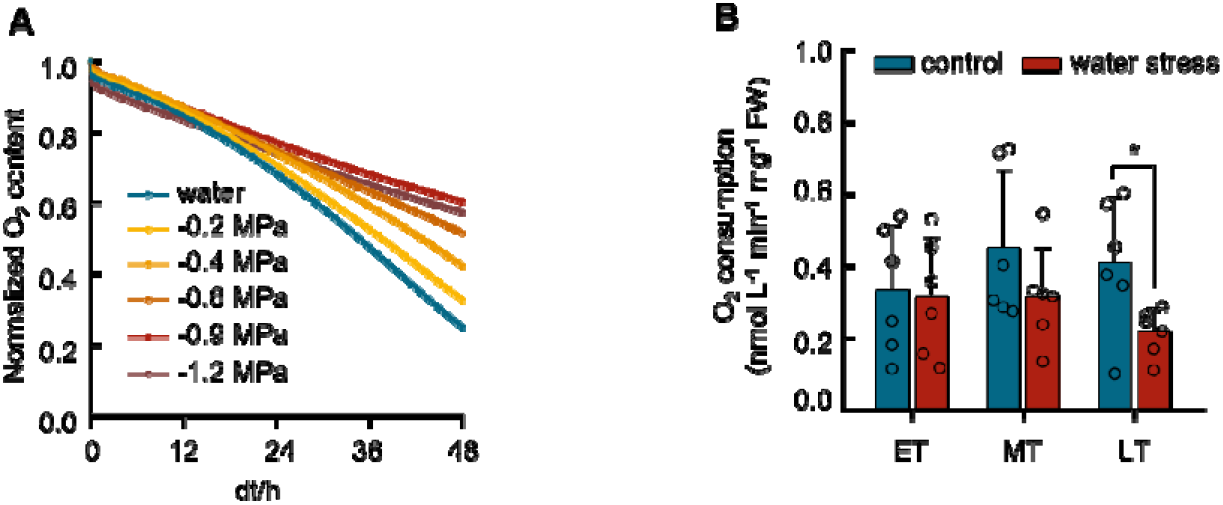
Water stress during germination alters seed respiration. (**A**) O_2_ consumption rate of stratified seeds imbibed on water or on a range of PEG solutions at 15°C in the dark. Data are means of 2 replicates of each concentration. (**B**) O2 consumption rate of stratified seeds imbibed at 3 timepoints of germination under control or water stress conditions (ET, early timepoint; MT, mid timepoint; LT, late timepoint). Data presented are means of 6 replicates ± SD. Student’s t-test was used to determine the statistical significance between the two groups at each time point. An asterisk below the bar chart indicates a significant difference between the two groups (*, P < 0.05).

The mitochondrial redox state was monitored during germination using the mitochondria-targeted roGFP2-Orp1 sensor with calibrated 405/488 nm excitation ratios as a proxy for redox potentials (Nietzel et al., 2019). To validate sensor functionality and determine its dynamic range in seeds, embryos were treated with 10 mM DTT and 100 mM H_2_O_2_ to obtain fully reduced and oxidized sensor states, respectively (Figure 4A, S1A). These treatments produced clear shifts in fluorescence ratios, confirming the probe’s responsiveness under the imaging conditions used (Figure 4B). In control conditions, the sensor oxidation level showed slight fluctuations during seed germination, suggesting the involvement of mitochondrial redox regulation in the germination process (Fig. 4A, B). A similar oxidation pattern was observed in seeds exposed to water stress (-0.9 MPa) throughout germination, although with a slight reduction in the sensor oxidation level (Figure 4B, C). We concluded that moderate water stress did not substantially alter the mitochondrial redox environment during germination, despite its inhibitory effect on germination kinetics and respiratory activity.

**Figure 4.**
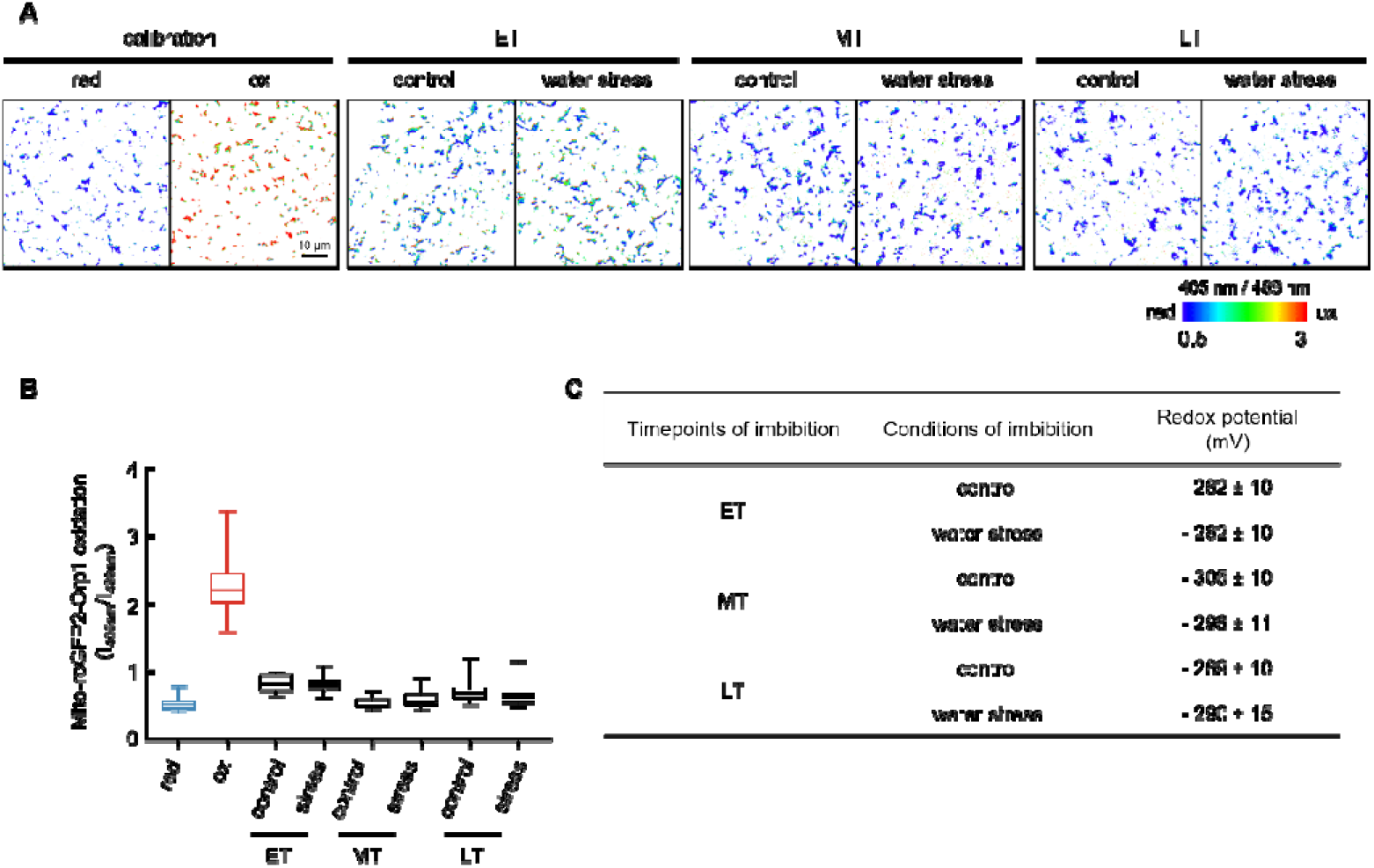
Mitochondrial roGFP2-Orp1 redox state during seed germination. (**A**) Confocal microscopy imaging of Mito-roGFP2-Orp1-emitted 405/488 fluorescence ratios in hypocotyl area upon seed imbibition in water (control) or a -0.9 MPa PEG solution (water stress) at 15°C in the dark. The Mito-roGFP2-Orp1 biosensor signal (left panel) was calibrated using 10 mM dithiothreitol (DTT) for full reduction or 100 mM H_2_O_2_ for full oxidation. (ET, early timepoint; MT, mid timepoint; LT, late timepoint) Scale bars = 10μm. (**B**) Redox ratio analysis at the three time points (**C**) Redox potentials at the three time points. In B and C, data were obtained from confocal imaging of at least 14 embryos in two independent experiments.

### Water stress perturbs ROS homeostasis

To further assess the involvement of ROS during germination under water stress, we used the ROS wheel model (Willems et al., 2016) to search for specific ROS footprints in our transcriptomic data (Table S4). The genes induced by water stress showed a significant overlap with Cluster III (“HL early”) following imbibition, indicating activation of an early redox signaling program typically associated with ROS-mediated stress responses and alterations in cellular oxygen homeostasis.

With increasing PEG concentrations, H_2_O_2_ production gradually decreased after 24 h of imbibition (Figure 5A). Consistently, H_2_O_2_ production increased progressively during germination under both control and water stress conditions (Figure 5B). However, water stress significantly decreased H_2_O_2_ production at all timepoints, resulting in approximately 1.3-and 1.5-fold lower levels at MT and LT, respectively, compared with the control. Likewise, catalase (CAT) activity also increased during germination but was significantly higher in PEG-treated seeds at the late stage (Figure 5C). Despite these changes, malondialdehyde (MDA) content remained unchanged between treatments at all time points (Figure 5D), indicating that water stress did not induce detectable oxidative damage to lipids. To further assess the contribution of ROS to germination under water stress, seeds were germinated in the presence of ROS scavengers. All scavengers had no effect on germination under control conditions, but an inhibitory effect was observed under water stress, except for DMTU, an H_2_O_2_ scavenger (Figure 5E). These results suggest that ROS production contributes positively to germination and that the reduced germination observed under water stress may be associated with altered ROS signaling.

To gain a spatial view of ROS production under water stress, we probed the fluorescent signal of H2DCFDA application. During the early and middle stages of germination, fluorescence signals appeared as discrete punctate structures distributed both within the cell and at the cell periphery (Figure 5F). As germination progressed, fluorescence became predominantly localized to intracellular punctate structures. Interestingly, fluorescence intensity appeared reduced under water stress conditions, particularly at the late stage of germination, consistent with the lower ROS accumulation observed in seed tissues. The punctate pattern of fluorescence suggests that ROS production is associated with specific subcellular compartments. Although H2DCFDA does not allow their precise identification, these structures may correspond to organelles involved in ROS metabolism, including peroxisomes, which are known to play important roles during seed germination and dormancy release (Pracharoenwattana et al., 2010; Jurdak et al., 2022). These observations indicate that ROS production is subject to a spatiotemporal regulation during germination and may be altered by water stress.

**Figure 5.**
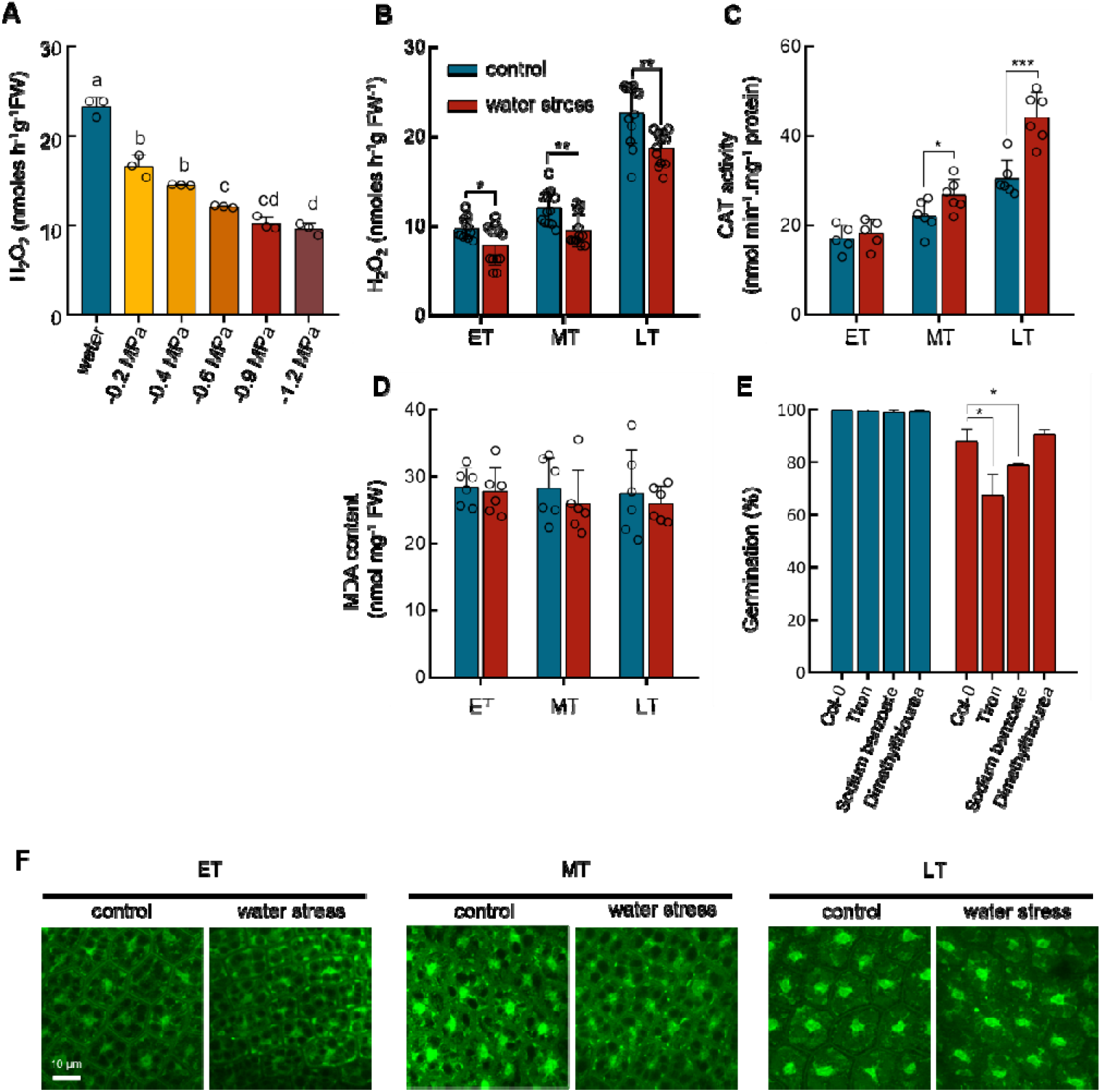
ROS homeostasis is involved in the response to water stress during germination. (**A**) H_2_O_2_ production of stratified seed imbibed with water or a range of PEG solutions for 24 h at 15°C under dark conditions. (**B**) H_2_O_2_ production (**C**) CAT activity and malondialdehyde (MDA) accumulation (**D**) at 3 time points during germination under normal or water stress conditions at 15°C in the dark. (**E**) Germination of stratified Col-0 seed after 10th days of imbibition at 15°C in the dark under normal or water stress conditions in the presence of ROS scavengers. Trion (0.1 mM; superoxide scavenger; Wise & Naylor, 1987); Sodium benzoate (0.1 mM; free radical scavenger; Hung & Kao, 2004); Dimethylthiourea (0.1 mM; Hydrogen peroxide scavenger; Levine & Lamb, 1994). (**F**) Localization of ROS production in embryonic axes of non-dormant seeds using 2_′_,7_′_-dichlorofluorescein diacetate (H2DCFDA) at 15 °C in the dark in the presence or absence of water stress. Data presented are means ± SD of three replicates (A and E), or 12 replicates (B and F), or six replicates (C and D). Student’s t-test was used to determine the statistical significance between treatments at each time point; an asterisk above the bar chart indicates a significant difference between the two groups (*, P < 0.05; **, P < 0.01; ***, P < 0.001).

To investigate a possible role of peroxisomes in ROS production, we monitored their redox status using the stably expressed peroxisome-targeted roGFP2-Orp1 transgenic reporter (Arnaud et al., 2023). Treatment of embryos with DTT and H_2_O_2_ led to significant differences in fluorescence ratios, thereby validating the sensor’s dynamic response to H_2_O_2_ (Figure S1B). Under control conditions, the redox level exhibited a slight increase at the middle time point, followed by a slight decrease at the late time point. However, these changes were not statistically significant. (Figure 6 A-C). Water stress significantly reduced peroxisomal sensor oxidation while preserving its overall temporal dynamics, with the strongest effect at the middle and late stages of germination (Figure 6A-C). Accordingly, the redox potential shifted to more negative values from −268 mV at ET to −299 mV at LT, indicating that peroxisomes remained more reduced under water stress throughout germination.

To confirm the role of peroxisomes in water stress, we assessed the expression of genes encoding peroxisomal glycolate oxidases, which produce H_2_O_2_ (Esser et al., 2014; Engqvist et al., 2015) and as *CAT2*, which encodes the major peroxisomal catalase responsible for H₂O₂ scavenging (Frugoli et al., 1996). These enzymes are therefore likely to contribute to the changes in peroxisomal redox status observed during germination under water stress. *GOX2* and *GOX3* transcript levels were lower under water stress than in control conditions (Figure S2A). Moreover, knock-out mutant lines for both genes are highly sensitive to water stress during germination (Figure S2C). In contrast, although *CAT2* transcript abundance was reduced under water stress, the *cat2* mutant did not display altered tolerance to water stress during germination (Figure S2B and S2D). Together, these findings indicate that peroxisomal ROS production contributes to the regulation of germination under water stress, and its lowering upon PEG treatment functionally contributes to seed germination delay.

**Figure 6.**
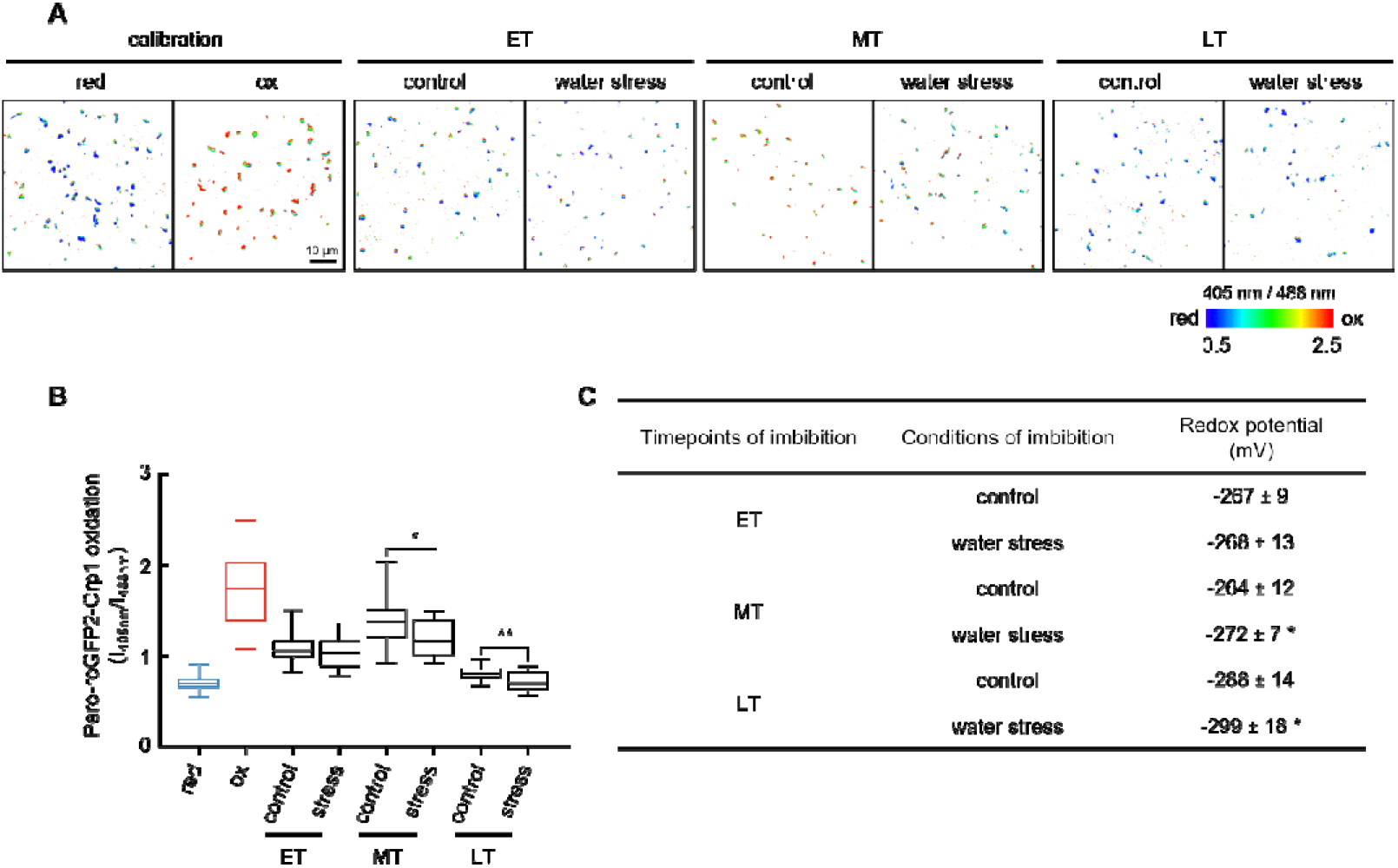
Peroxisomal redox status is altered by water stress during germination. (**A**) Confocal microscopy imaging of 405/488 sensor redox ratios depicted as false-colour scales in hypocotyls upon seed imbibition with water (control) or -0.9 MPa PEG solution (water stress) at 15°C in the dark. RoGFP2-Orp1 (left panel) was calibrated using 10 mM dithiothreitol (DTT) for full reduction (Red) or 100 mM H2O2 for full oxidation (Ox) of the biosensor in vivo. (ET, early timepoint; MT, mid timepoint; LT, late timepoint) Scale bars = 10μm. (**B**) RoGFP2-Orp1Roxydation levels at the three time points. (**C**) Redox potentials calculated from confocal microscopy images. In B and C, data were obtained from at least 14 embryos in two independent experiments. Statistical significance was determined using Student’s t-test at each time point (*, P < 0.05; **, P < 0.01).

### ROS-associated gene networks and hormonal pathways regulate germination under water stress

To identify the key regulatory genes associated with H₂O₂ production, we performed a weighted gene co-expression network analysis (WGCNA) integrating our transcriptomic and H₂O₂ production data (Figures 7A and S3). A soft-thresholding power of β = 6 was selected to construct the co-expression network, resulting in the identification of 31 co-expression modules (Figures S3A, S3B and 7A). Among these, the purple module, comprising 335 genes, showed the strongest positive correlation with ROS production (Figure 7A). The eigengene of this module exhibited distinct temporal expression changes during germination (Figure S3C), indicating coordinated regulation of its constituent genes. Gene Ontology enrichment analysis revealed a significant overrepresentation of biological processes related to cell wall organization, external encapsulating structure organization, transmembrane transport, and negative regulation of abscisic acid (ABA)-activated signaling (Figure 7B), suggesting a close association between this module and H₂O₂ production. We further visualized the genes within the purple module using Cytoscape and identified five highly connected hub genes, including AT1G20440 (*COLD-REGULATED 47*, *COR47*), AT1G73220 (*ORGANIC CATION TRANSPORTER 1*, *OCT1*), AT1G28290 (*ARABINOGALACTAN PROTEIN 31*, *AGP31*), and two genes with no annotated function, AT4G11100 and AT3G15940 (Figure 7C). The module also contained several transcription factor genes, including members of the MYB, C2H2 zinc-finger, NAC, and GRF families (Table S5), suggesting that transcriptional regulation contributes to the coordination of ROS-associated responses during germination.

**Figure 7.**
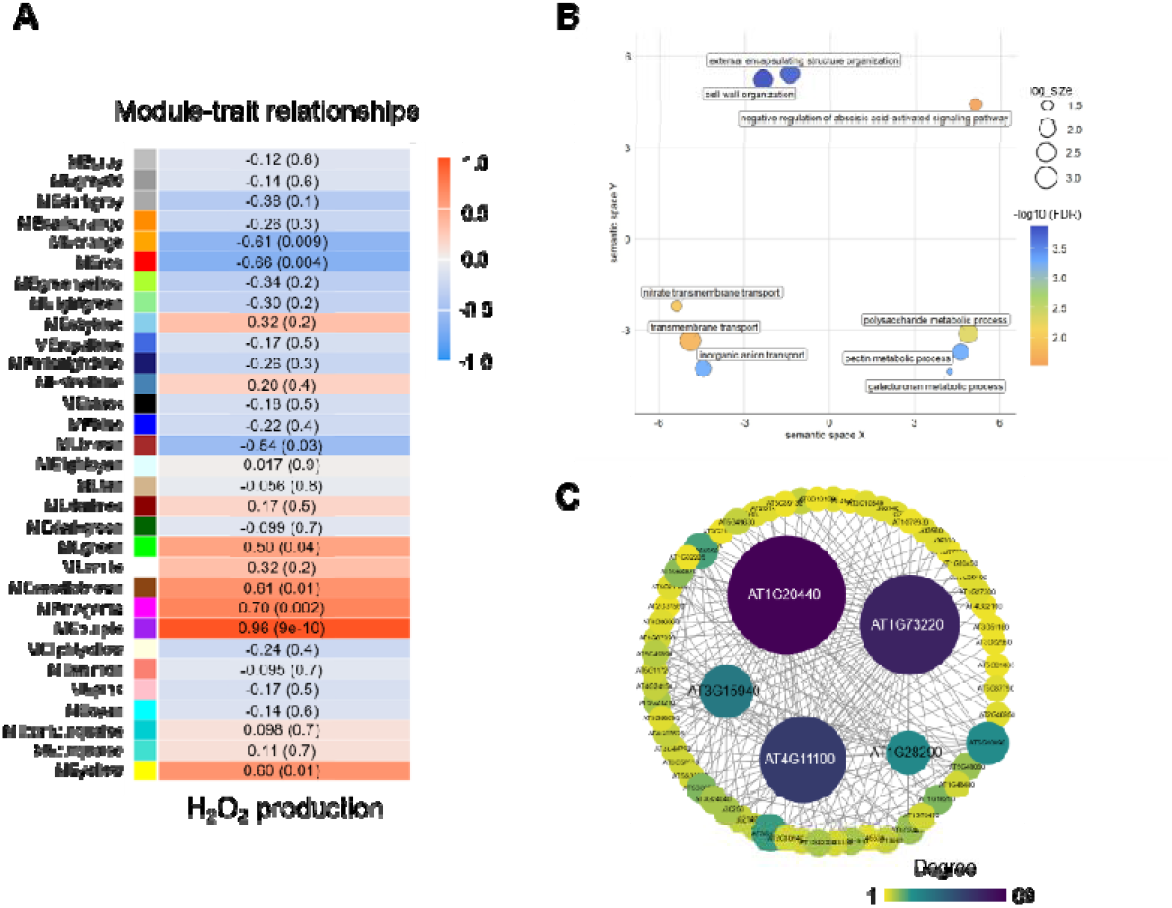
A central gene hub is associated with germination and ROS production under water stress. (**A**) Correlation matrix between WGCNA modules and ROS production under water stress conditions. The x-axis represents ROS production, the y-axis represents the name of each module, the P value is presented in each cell, and the color of each cell indicates the correlation between the module and the ROS production. (**B**) Simplified gene ontology classification of the purple module . (**C**) Correlation network visualization of the interactions between the five genes with highest connectivity in the purple module. Each node is labelled with its corresponding gene IDs. Node size represents the degree of correlation among genes.

We examined the transcriptomic data to gain deeper information on the possible role of the ABA pathway in ROS-mediated responses to water stress during germination (Figure 8A). Consistent with the WGCNA analysis, water stress markedly altered the expression of genes associated with ABA signaling (Figure 8B). In particular, *SnRK2.2* and *SnRK2.5*, which encode protein kinases required for ABA signaling (Fujii et al., 2007; Wang et al., 2015), were strongly induced under water stress conditions. In addition, expression of the ABA-responsive transcription factor ABF3, whose activity depends on ABA signaling (Yoshida et al., 2010), was significantly upregulated, both indicating an activation of the ABA signaling pathway in response to water stress during germination. Several genes encoding key enzymes of the ABA biosynthetic pathway, including *NCED2* (9-cis-epoxycarotenoid dioxygenase 2; Zhang et al., 2025) and *ABA2* (short-chain dehydrogenase/reductase; González-Guzmán et al., 2002), were significantly upregulated by water stress (Figure 8B). Conversely, *CYP707A2* and *CYP707A3*, which encode ABA 8′-hydroxylases involved in ABA catabolism (Saito et al., 2004; Kushiro et al., 2004), were significantly downregulated under water-stress conditions (Figure 8). To assess the role of ABA in water-stress-induced germination inhibition, seeds were treated with the ABA biosynthesis inhibitor fluridone (Yoshioka et al., 1998). Surprisingly, inhibition of ABA biosynthesis did not alleviate the inhibitory effect of water stress on germination (Figure 8C). In contrast, genetic analyses supported a role for ABA in this response. Germination analysis of the *nced2,5,6,9* quadruple mutant line affected in ABA biosynthesis or the *abi4* ABA signaling mutant line showed that ABA is required for water-stress-induced germination inhibition (Figure 8D). Remarkably, seeds of these mutant lines also exhibited significantly higher ROS accumulation than wild-type seeds under water stress conditions (Figure 8E). Collectively, these results demonstrate that ABA signaling contributes to the inhibition of germination under water stress and unravel a functional link between ABA and ROS pathways.

**Figure 8.**
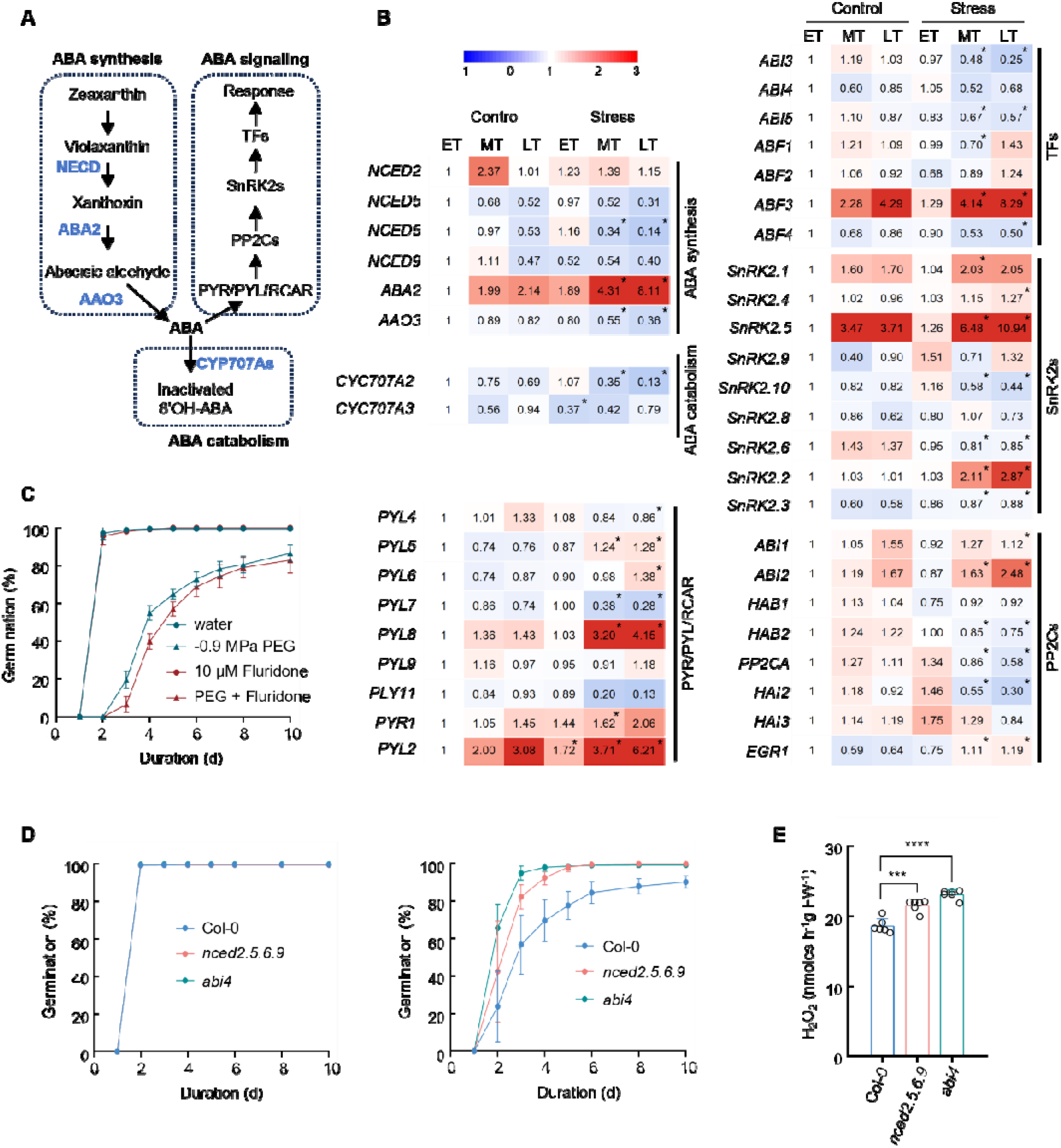
ABA pathways respond to water stress during germination. (**A**) Schematic representation of the ABA pathway, including synthesis, catabolism, and signaling. (**B**) Heat maps showing differentially-expressed genes in the ABA pathway in seeds exposed to water (control) or PEG at -0.9 MPa. For each gene, transcript abundance is expressed as normalized read counts relative to the early time point (ET) in the control condition, which was arbitrarily set to 1. Asterisks indicate significant differences relative to the corresponding control condition at the same time point (FDR < 0.01). (**C**) Germination of stratified seeds at 15°C in the dark under normal or water stress conditions in the presence of 10 µM fluridone. (**D**) Germination at 15°C in the dark under normal (left) or water stress (right) conditions of seeds of ABA pathway-associated mutants. (**E**) H2O2 production of ABA-associated mutants at 15°C in the dark under water stress conditions after 48h of imbibition (LT). Data presented are means ± SD of three replicates (B, C, and D) or six replicates (E). Student’s t-test was used to determine the statistical significance between two lines; an asterisk above the bar chart indicates a significant difference between the two groups (***, P < 0.001; ****, P < 0.0001).

Considering that the antagonistic interplay between ABA and GA is a central determinant of the balance between dormancy maintenance and germination completion, we also examined the expression of genes involved in GA biosynthesis, catabolism, and signaling (Figure 9A and 9B). The *GA3ox2* gene, encoding a key enzyme for GA biosynthesis (Mitchum et al., 2006), was strongly downregulated under water stress (Figures 9A and 9B). Conversely, the *GA2ox6* and *GA2ox8* genes, which encode GA 2-oxidases involved in GA catabolism (Kubalová et al., 2025), were significantly upregulated. These observations suggest that water stress reduces the pathways leading to bioactive GA accumulation during germination. Consistently, water stress also affected the expression of GA signaling genes, notably *GID1* encoding the GA receptor (Olszewski et al., 2002) (Figure 9A and 9B). To assess the role of GA in the germination response to water stress, seeds were treated with paclobutrazol (Pac), an inhibitor of GA biosynthesis. Pac treatment significantly exacerbated the inhibitory effect of water stress on germination (Figure 9C), supporting a positive role for GA in promoting germination under low water potential. Interestingly, Pac treatment also triggered a significant decrease in ROS accumulation (Figure 9D), suggesting a functional link between GA signaling and ROS homeostasis during germination.

**Figure 9.**
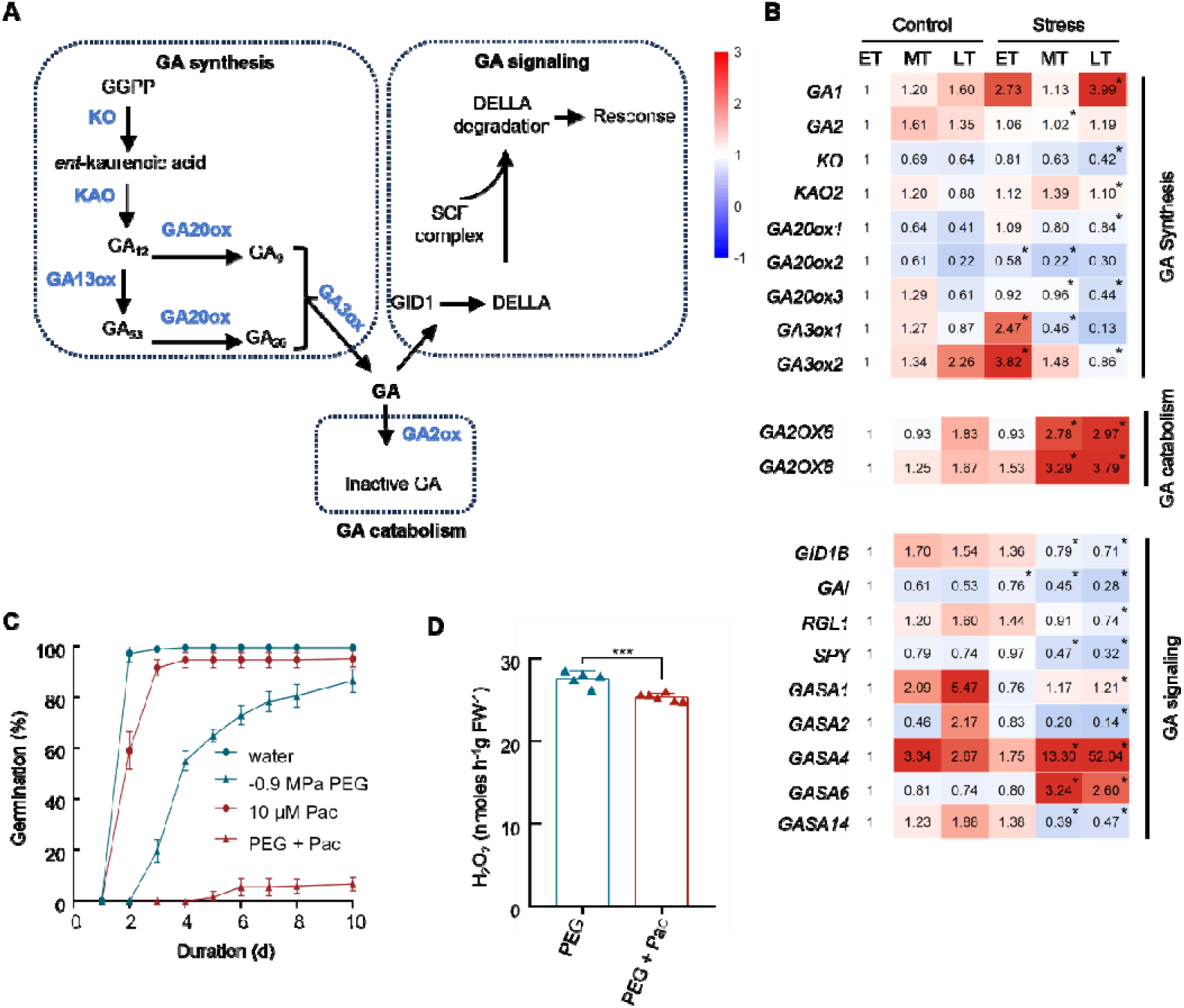
GA pathways respond to water stress during germination. (**A**) Schematic representation of the GA pathway, including synthesis, catabolism, and signaling. (**B**) Heat maps showing differentially-expressed genes associated with GA metabolism and signaling pathways in seeds exposed to water (control) or PEG at -0.9 MPa. For each gene, transcript abundance is expressed as normalized read counts relative to the early time point (ET in the control condition, which was arbitrarily set to 1. Asterisks indicate significant differences relative to the corresponding control condition at the same time point (FDR < 0.01). (**C**) Germination assay of seeds treated with GA inhibitor (Paclobutrazol, Pac) under control or water-stress conditions. (**D**) H_2_O_2_ production in non-dormant Col seeds imbibed with -0.9 MPa PEG for 48 h (LT) in the presence or absence of Paclobutrazol. Data are presented as means ± SD of 3 replicates (C) or at least 5 replicates (D). Student’s t-test was used to determine the statistical significance between two lines; an asterisk above the bar chart indicates a significant difference between the two groups (***, P < 0.001).

## DISCUSSION

### Water stress induces a developmental slowdown rather than a metabolic arrest

Decreasing water potential to -0.9 MPa delayed germination without substantially affecting its completion, indicating that seeds retained the capacity to germinate (Figure 1A). Similar effects have been reported in several plant species (Vallejo et al., 2010; Saux et al., 2020; Wang et al., 2022; Ouzakar et al., 2026). Importantly, the moderate reduction in seed water content observed at -0.9 MPa indicates that delayed germination cannot be explained solely by impaired hydration. Instead, the progressive increase in the number of differentially expressed genes throughout germination (Figure 2A) indicates that water stress triggers a continuous transcriptional reprogramming rather than a transient response during imbibition. Thus, moderate water stress temporarily restrains developmental progression, which delays radicle protrusion while preserving germination competence until conditions become more favorable.

Consistently, water-stress-induced transcriptome dynamics were found to be highly selective rather than indicative of a generalized metabolic suppression. Consistent with our observations of H_2_O_2_ accumulation under optimal germination conditions, genes associated with oxygen responses, redox metabolism, signaling, and stress adaptation were induced in response to water stress. In contrast, genes involved in RNA processing, ribosome biogenesis, DNA replication, and chromatin organization were strongly downregulated (Figure 2). Related transcriptomic reprogramming in response to water limitation during germination has been reported in seeds of lettuce (Hwang et al., 2024), *Allium mogolicum* (Men et al., 2026), and sunflower (Saux et al., 2020). This pattern suggests that water stress responses actively prioritize stress adaptation over growth, a deceleration strategy also noted in drought-exposed vegetative tissues, where growth is rapidly restricted while protective pathways remain active (Skirycz and Inzé, 2010; Wilkins et al., 2010). Here, as discussed below, we identified a temporal regulatory framework linking water stress perception to delayed germination through coordinated events that unfold during seed imbibition.

### Water stress rapidly reallocates cellular resources from germination-associated growth to stress-adaptation processes before major physiological inhibition occurs

A striking observation of this study is that the transcriptional response to PEG occurred before the appearance of major physiological differences. For instance, at ET, relatively few genes were differentially expressed, yet their functional identities provide important insights into the initial perception of water stress and specific stress signaling pathways, altogether showing that seeds rapidly activate protective mechanisms in response to reduced water availability. Concomitantly, genes associated with cell wall metabolism and xyloglucan remodeling were repressed. Because cell wall loosening and endosperm weakening are prerequisites for radicle emergence (Finch-Savage and Leubner-Metzger, 2006; Steinbrecher and Leubner-Metzger, 2017), repression of these pathways suggests that water stress affects germination from its earliest stages by limiting the acquisition of growth competence. Interestingly, sulfur metabolism was among the earliest pathways to be activated. Sulfur assimilation supplies cysteine for glutathione biosynthesis and is therefore closely linked to cellular antioxidant capacity (Kopriva and Rennenberg, 2004). Its induction suggests that redox homeostasis is adjusted early during germination, before measurable changes in H_2_O_2_ accumulation become apparent. Together, these observations reveal that perception of water deficit by germinating seeds rapidly redirects transcriptional and downstream resources toward stress adaptation while postponing developmental processes required for embryo expansion and radicle protrusion.

### Water stress reduces germination-associated H_2_O_2_ accumulation while activating redox-sensitive pathways

ROS have emerged as central regulators of seed dormancy release and germination (Bailly, 2004, 2019, 2022). According to the “oxidative window” concept, successful germination requires ROS to accumulate within an optimal range that promotes signaling while avoiding oxidative damage (Bailly et al., 2008). Our data indicate that the transition from the early to the middle stage of germination represents a critical checkpoint at which water stress progressively shifts seeds away from this permissive redox state. At the middle stage, PEG treatment induced extensive transcriptional reprogramming dominated by oxygen-and redox-related processes, including responses to hypoxia, oxygen deprivation, electron transport, oxidative phosphorylation, and redox metabolism. Consistent with this, PEG-responsive genes were significantly enriched within the high-light (HL) ROS cluster defined by Willems *et al*. (2016), indicating activation of a coordinated redox-responsive transcriptional program. However, physiological analyses revealed that this transcriptional signature is not reflected by enhanced ROS accumulation. Instead, PEG-treated seeds accumulated less H₂O₂, exhibited higher catalase activity, and showed no increase in lipid peroxidation, demonstrating that moderate water stress does not induce oxidative damage but rather restricts the accumulation of signaling ROS. This interpretation is further supported by the inhibitory effect of ROS scavengers on germination, confirming that ROS are required for germination progression. These findings illustrate that transcriptomic signatures associated with oxidative stress should not be interpreted as direct evidence of increased ROS production but instead reflect activation of adaptive responses to altered redox homeostasis.

Subcellular imaging of redox status revealed that this response was highly compartmentalized. Whereas the mitochondrial redox potential remained remarkably stable despite reduced respiration, the oxidative status of peroxisomes decreased markedly under water stress. Given the established roles of peroxisomes in reserve mobilization, β-oxidation, dormancy release, and H_2_O_2_ production during germination (Oracz et al., 2007; Pracharoenwattana et al., 2010; Jurdak et al., 2022), these results identify peroxisomes as a major site of redox regulation under water deficit. We propose that moderate water stress imposes a reversible growth restraint by limiting the accumulation of peroxisome-derived signaling H_2_O_2_ to levels below the threshold required to sustain the developmental transitions leading to radicle protrusion, rather than by triggering a damaging oxidative burst.

This mechanism contrasts with the increased ROS production typically observed in drought-stressed vegetative tissues (Liu et al., 2019; Wang et al., 2022), highlighting the importance of considering developmental context when interpreting plant redox responses. More broadly, our findings reinforce the concept that ROS function primarily as developmental signals during germination and demonstrate that successful adaptation to water stress relies on maintaining ROS within an appropriate signaling window rather than maximizing antioxidant protection. They also emphasize the importance of integrating physiological measurements with transcriptomic analyses to accurately resolve the mechanisms underlying stress responses during seed germination.

### Peroxisomes emerge as a central redox signaling hub controlling germination under water stress

The compartment-specific redox analyses identify peroxisomes as a major redox signaling hub linking water stress to delayed germination. While peroxisomes have long been associated with reserve mobilization, dormancy release, and germination (Oracz et al., 2007; Leymarie et al., 2012; Jurdak et al., 2022), our transcriptomic analyses indicate that water stress selectively reprograms peroxisomal functions rather than globally suppressing it. Because acyl-CoA oxidases constitute a major source of peroxisomal H_2_O_2_ during germination (Eastmond et al., 2000; del Río et al., 2006), the downregulation of key genes involved in peroxisomal H_2_O_2_ production provides a molecular explanation for both the reduced H_2_O_2_ accumulation and the more reduced peroxisomal redox state observed under water stress. Interestingly, this response was not associated with a complete shutdown of peroxisomal metabolism. In particular, genes encoding acyl-CoA oxidases, which constitute the major source of peroxisomal H_2_O_2_ during germination (Eastmond et al., 2000; del Río et al., 2006), together with *PEX5*, were repressed under PEG treatment, providing a molecular explanation for the reduced peroxisomal oxidation observed in vivo. In contrast, genes involved in downstream β-oxidation and the glyoxylate cycle, including 3-ketoacyl-CoA thiolase-2 (*KAT2,* Germain et al., 2001) and Isocitrate Lyase (*ICL*, Graham, 2008), remained expressed, suggesting that carbon mobilization is maintained while ROS-production is selectively attenuated.

The functional significance of this regulation is supported by the enhanced sensitivity of *gox1*, *gox2*, and *gox3* mutant seeds to water stress, demonstrating that impairment of peroxisomal H_2_O_2_ production compromises germination under low water potential. By contrast, the absence of a germination phenotype in the *cat2* mutant indicates that reducing peroxisomal H_2_O_2_ scavenging alone is insufficient to modify the response to water stress. Together with the transcriptomic and redox imaging data, these genetic analyses indicate that the production of peroxisome-derived H_2_O_2_, rather than its detoxification, is a critical determinant of germination progression under water deficit. We therefore propose that water stress selectively suppresses a permissive peroxisomal H_2_O_2_ signal while preserving the metabolic functions required for reserve mobilization and seedling establishment.

### WGCNA identifies developmental and hormonal pathways associated with peroxisomal H_2_O_2_ signaling

The WGCNA analysis provided a systems-level perspective on the biological significance of the altered peroxisomal redox state observed under water stress. Notably, one of the modules showing the strongest correlation with H_2_O_2_ accumulation was enriched in genes involved in cell wall organization and hormonal regulation. The strong coregulation between H_2_O_2_-related modules and genes involved in cell wall organization suggests that reduced peroxisomal H_2_O_2_ under water stress delays germination by impairing the cell wall remodeling processes required for radicle protrusion. The identification of ABA-related genes within a H_2_O_2_-associated gene hub, together with the strong correlation between H_2_O_2_ accumulation and developmental processes required for germination, suggests that altered redox signaling may represent an upstream event contributing to the hormonal reprogramming induced by PEG. This is consistent with the extensive crosstalk between H_2_O_2_, ABA, and GA signaling pathways during seed germination (Liu et al., 2010; Ishibashi et al., 2012; Bi et al., 2017; Cheng et al., 2022) and raises the possibility that suppression of peroxisome-derived H_2_O_2_ constitutes one of the primary signals through which water stress modifies hormonal balance and developmental timing.

Increasing evidence indicates that H_2_O_2_ functions both upstream and downstream of ABA and GA signaling, forming an interconnected regulatory network that controls seed germination (Bailly, 2008, 2022; Liu et al., 2010). Our results support the existence of such a regulatory network under water stress. Transcriptomic analyses revealed activation of ABA biosynthesis and signaling together with repression of ABA catabolic genes. Consistent with these observations, both ABA-deficient and ABA-insensitive mutants exhibited a reduced germination delay and accumulated higher levels of H_2_O_2_ under water stress (Figure 8D), indicating that ABA contributes to limiting H_2_O_2_ accumulation during germination. This concurs with previous studies showing that ABA modulates cellular redox homeostasis through the regulation of both antioxidant systems and ROS-producing pathways. Conversely, water stress repressed the expression of genes associated with GA biosynthesis and signaling. Moreover, pharmacological inhibition of GA biosynthesis further reduced H_2_O_2_ accumulation, supporting previous reports that GA promotes germination, at least in part, by stimulating ROS production (Müller et al., 2009; Bahin et al., 2011). Together, these findings indicate that ABA and GA exert opposite effects on ROS homeostasis during germination: ABA restricts, whereas GA promotes, the accumulation of signaling H_2_O_2_.

Based on our physiological, genetic, and transcriptomic data, we propose that peroxisome-derived H_2_O_2_ constitutes a central regulatory hub coordinating ABA and GA interplay under water stress. Moderate water deficit initially reduces peroxisomal H_2_O_2_ production, thereby shifting the hormonal balance toward enhanced ABA signaling and reduced GA responsiveness. These hormonal changes further reinforce the decline in H_2_O_2_ accumulation, establishing a positive feedback loop that progressively stabilizes a low-redox state. As a consequence, downstream processes required for germination, including cell wall remodeling, embryo growth, and ultimately radicle protrusion, are delayed. Rather than acting as independent signaling pathways, ABA, GA, and H_2_O_2_ therefore appear to constitute an integrated regulatory network organized around peroxisomal redox signaling.

### Water stress maintains a transcriptionally and translationally restrained developmental state

Our transcriptomic analyses indicated that water stress progressively triggers a transcriptionally and translationally restrained developmental state during germination. This response was pronounced at mid- and late-germination time points, where the most significantly enriched downregulated GO categories were associated with ribosome biogenesis, rRNA processing, RNA metabolism, ribonucleoprotein complex assembly, DNA replication, chromosome organization, and ATP-dependent chromatin remodeling. Given the downregulation of multiple genes encoding ribosomal and nucleolar proteins, water stress may, therefore, limit the capacity of germinating seeds to sustain the increase in translational activity required for embryo growth, thereby contributing to delayed germination under water stress.

Importantly, these regulatory mechanisms were accompanied by a broad downregulation of genes involved in chromatin assembly, transcriptional regulation, RNA processing, and genome organization, suggesting that water stress limits the chromatin reconfiguration normally associated with developmental progression. Chromatin modification and remodeling are increasingly recognized as critical components of dormancy release and germination (Nonogaki, 2019; Sano et al., 2022), and the widespread repression of chromatin-associated functions observed during germination suggests that PEG-treated seeds fail to fully activate the transcriptional programs required for rapid growth.

Finally, these changes did not reflect developmental arrest. Genes associated with chloroplast biogenesis and photosynthesis were efficiently induced under water stress, indicating that developmental progression continued despite its delay. Instead, the downregulation of translational and chromatin-remodeling pathways suggests that water stress maintains seeds in a state of reduced developmental competence, thereby delaying the transition toward rapid embryo growth and radicle protrusion.

Altogether, we provided evidence that delayed germination under water stress results from a coordinated developmental reprogramming rather than a simple consequence of reduced water availability, and we identified a temporal sequence of events linking water-stress perception to delayed radicle protrusion. It involves an initial activation of stress-acclimation responses, and subsequent reduction in peroxisome-derived ROS accumulation, the promotion of ABA dominance over GA responsiveness, and plausibly, the establishment of a transcriptionally and translationally restrained developmental state that contributes to germination delay. These data support a model in which water stress primarily uncouples the metabolic and signaling functions of peroxisomes, maintaining reserve mobilization while attenuating ROS-dependent developmental signaling. This altered redox state propagates through hormonal and developmental networks controlling cell wall remodeling, embryo growth, and germination timing. Together, these findings provide a mechanistic framework explaining how seeds dynamically adjust germination to water availability and identify peroxisome-derived ROS as a key regulator of environmental control of germination.

## MATERIALS AND METHODS

### Plant material

*Arabidopsis thaliana* seeds used in this study were from the Columbia (Col-0) genetic background. All mutant lines and reduction–oxidation sensitive Green Fluorescent Protein 2 fused to Oxidant Receptor Peroxidase 1 (roGFP2-Orp1) lines (Nietzel et al., 2019) were derived from the same background. We used the ABA related mutants, nced2.5.6.9 quadruple mutant (Chauffour et al., 2019) and abi4 single mutant (Dong et al., 2016), glycolate oxidase mutants (gox1-2, gox2-2, and gox3-2; Yang et al., 2025) and catalase mutant (cat2-2; Yang et al., 2025). Prior to sowing, seeds were stratified at 4°C for 4 days, then planted in soil and transferred to growth chambers maintained under controlled conditions (21°C, 16-hour daily photoperiod, and light intensity of 150 µmol m^-2^ s^-1^). Once matured, seeds were collected and stored at -20°C until required for further experiments.

### Seed germination assays

Seeds were stratified prior to germination assays by placing them on a layer of cotton wool moistened with deionized water in darkness at 4 °C for 4 days. For germination assays, stratified seeds were transferred to petri dishes containing two layers of cotton overlaid with filter paper, moistened with either distilled water or polyethylene glycol 8000 (PEG 8000) solutions. A range of PEG solutions was prepared to achieve water potential values from -0.2 to -1.2 MPa, as described by Michel and Kaufmann (1973), depending on experimental requirements. The concentrations of the chemical reagents used for germination are provided in the main text. All assays were conducted in the dark and included at least 50 seeds per genotype or treatment with a minimum of 3 biological replicates. Germination was monitored daily for 10 days. A seed was considered germinated when the radicle had emerged through the seed coat.

### Seed moisture content

Approximately 10 mg of *Arabidopsis* seeds were imbibed with water (control) or -0.9 MPa PEG (water stress) at 15°C in darkness. At the indicated time points, seeds were collected and immediately weighed to determine fresh weight (FW). The collected samples were then dried at 100°C for 24 h to determine dry weight (DW). Water content was calculated and expressed as g H_2_O g^-1^ FW. Data represent the mean ± SD of three biological replicates.

### Measurement of hydrogen peroxide production

Hydrogen peroxide production was assessed by a decrease in scopoletin fluorescence, as described by Leymarie et al. (2012). Twenty-four mg of seeds were incubated in 200 µl of potassium phosphate buffer (20 mM, pH 6.0) containing 5 µM scopoletin and 1 U/mL of horseradish peroxidase (HRP) in darkness at 25°C on a shaker for 1 h, and known H_2_O_2_ concentrations were used for calibration. Measurements were carried out at excitation/emission wavelengths of 346 nm/455 nm. Results correspond to the mean ± SD of 3 replicates and are expressed per mg of the fresh weight.

### Malondialdehyde (MDA) content

Malondialdehyde (MDA) was assessed as described by Heath and Packer (1968). Approximately twelve mg of *Arabidopsis* seeds were homogenized in 0.1% trichloroacetic acid (TCA) on ice. Following centrifugation at 12,000 × g for 10 min, the supernatant was collected and mixed with an equal volume of 0.5% thiobarbituric acid (TBA). The reaction mixture was incubated at 95°C for 30 min, then cooled on ice for 10 min to stop the reaction. Absorbance was measured at 532 and 600 nm using a spectrophotometer. MDA content was determined using an extinction coefficient of 155 mM^-1^ cm^-1^. Results correspond to the mean ± SD of 3 replicates and are expressed per g of the fresh weight.

### Measurement of catalase activity

Approximately fifty mg of *Arabidopsis* seed were ground with liquid nitrogen, and proteins were extracted in potassium phosphate buffer (0.1 M, pH 7.8) containing 2 mM dithiotreitol, 0.1 mM EDTA, and 1.25 mM polyethylene glycol 4000 as described by Bailly et al (1996). After centrifugation at 13000 g for 15 min, the supernatant of each sample was desalted on Sephadex PD10 columns and used for measuring catalase (CAT) activity. CAT activity is expressed as nanomole H_2_O_2_ per mg per protein per min, and data presented are means ± SD.

### *In situ* localization of ROS

Intracellular localization of ROS was visualized using 2’,7’-dichlorodihydrofluorescein diacetate (H2DCFDA) with a confocal microscope as described by Leymarie et al (2012). Embryos were dissected from imbibed seeds and incubated in 50 mM potassium phosphate buffer (pH 7.4) containing 25 µM H2DCFDA for 30 min at 20 °C in the dark. Following incubation, embryos were rinsed in potassium phosphate buffer for 10 min. Images were acquired using a confocal laser scanning microscope (Leica TCS SP5 AOBS) with excitation at 488 nm and emission collected between 498 and 550 nm, using a 63 X objective. Z-stack images were collected with a step size of 0.5 µm. To compare H2DCFDA fluorescence intensity, embryonic axes from different experimental conditions were fixed simultaneously and imaged using identical confocal settings. The acquired images were further processed using ImageJ software.

### Confocal imaging of roGFP2-Orp1 lines

Confocal imaging of imbibed *Arabidopsis thaliana* seed expressing roGFP2-Orp1 was acquired using a confocal laser scanning microscope (Zeiss LSM 980 inverted) equipped with a Plan-Apochromat 63X/1.40 water-immersion objective. For calibration, de-coated seeds were fully reduced or oxidized by treatment with 10 mM dithiothreitol (DTT) or 10 mM H_2_O_2_, respectively. roGFP2-Orp1 was sequentially excited at 405 nm and 488 nm, and fluorescence emission was collected between 508 and 530 nm. The acquired images were visualized, and fluorescence quantified using Matlab-based software (Fricker, 2016), and the redox potential was calculated according to the method described by (Schwarzländer et al., 2008).

### Respiration measurement

Approximately 5 mg of seeds were stratified at 4°C for 4 days and then transferred to 2 mL glass vials containing cotton pads saturated with either distilled water or PEG solution (-0.9 MPa). Oxygen content in the vials was monitored using a fixed-needle oxygen microsensor (PyroScience; OXF500PT-OI) calibrated with air and nitrogen before measurement. Oxygen content was recorded every 10 min for 2 days with FireStingGO2 Manager software connected to the sensor. Respiration rate was defined as the change in oxygen uptake within 10 min intervals before and after the selected time point. Data correspond to the mean ± SD of six replicates and are expressed as nmol L-1 of oxygen consumed per min per mg of fresh weight.

### RNA isolation and transcriptomic analysis

Total RNA was extracted from approximately 50 mg of seeds using the cetyltrimethylammonium bromide (CTAB) method as described by Chang et al (1993). Genomic DNA of the sample was removed using the TURBO DNA-free™ Kit and purified using the NucleoSpin™ RNA Clean-up XS kit. RNA samples with RNA concentration > 100 ng/µL and RNA integrity number (RIN) values > 4.0 were deemed suitable for further sequencing.

Library construction and subsequent sequencing were performed on the Novogene Illumina platform. Sequencing was performed using paired-end sequencing, with each end containing 150 bp reads (PE150); approximately 6 Gb of data per sample was acquired. After sequencing, raw reads were quality-filtered and aligned to the *Arabidopsis* reference genome (Araport11 annotation of the TAIR10 genome) using Hisat2 v2.0.5. Differential expression analysis was conducted using the DESeq2 package (v1.20.0) in R software. Genes with a false discovery rate (FDR) < 0.01 were considered significantly differentially expressed under water stress conditions. Gene set enrichment analysis (GSEA) was used to assess the enrichment of differentially expressed genes in the Gene Ontology (GO) and Kyoto Encyclopedia of Genes and Genomes (KEGG) databases. Significantly enriched pathways were identified based on normalized enrichment scores (NES) and corrected p-values.

### Weighted gene co-expression network analysis

To identify hub genes associated with ROS generation, weighted gene co-expression network analysis (WGCNA) was performed using the WGCNA R software package. Genes with expression levels below 1 in more than 50% of the samples were excluded. The filtered genes were used to determine the soft-thresholding power (β) using the pickSoftThreshold function, with the optimal value selected when the scale-free topology fit index first exceeded 0.9. Using the selected soft-thresholding power (β = 6, Figure S3A), the gene expression data were clustered into 31 modules based on their co-expression patterns. Module eigengenes were then calculated, and their correlations with ROS generation (including control and water stress conditions) were assessed using Pearson correlation analysis. A module with an FDR < 0.01 and an absolute correlation coefficient (|r|) > 0.75 was considered significantly associated with ROS production. Gene co-expression networks within the selected module were visualized using Cytoscape software (v3.10.2). Genes with the top five degree values were defined as hub genes for further analysis.

### Statistical analyses

Statistical analyses were performed using Student’s t-test for comparisons between two groups and analysis of variance (ANOVA) for comparisons among three or more groups with GraphPad Prism. Differences were considered statistically significant at *P* < 0.05.

## DATA AVAILIBILITY STATEMENT

The RNA-seq data generated in this study have been have been deposited in the NCBI Gene Expression Omnibus (GEO) under the GSE338026 accession.

## ACKNOWLEDGEMENTS

This work benefited from a China Scholarship Council (CSC) doctoral scholarship to XM, the research grant SEED (ANR-25-CE20-6227-02) from Agence Nationale pour la Recherche (ANR) to CB and FB. F.B. was also supported by Velux Stiftung (project No. 1747). The authors thank Pr Yi Han (China Agricultural University) for providing the Arabidopsis *gox1-2*, *gox2-2*, *gox3-2*, and *cat2-2* mutant seeds, and Dr Annie Marion-Poll (INRAe) for providing the Arabidopsis *nced2 nced5 nced6 nced9* quadruple mutant and *abi4* mutant seeds.

## AUTHOR CONTRIBUTIONS

CB and FB conceived and designed the research. XM performed the experiments and analyzed the data. AV contributed to the transcriptome data analysis. SW provided technical assistance to XM. XM wrote the original draft of the manuscript. CB, FB, and AM revised the manuscript. All authors read and approved the final manuscript.

## SUPPORTING INFORMATION

The following Additional Supporting Information is available for this article:

**Figure S1.**
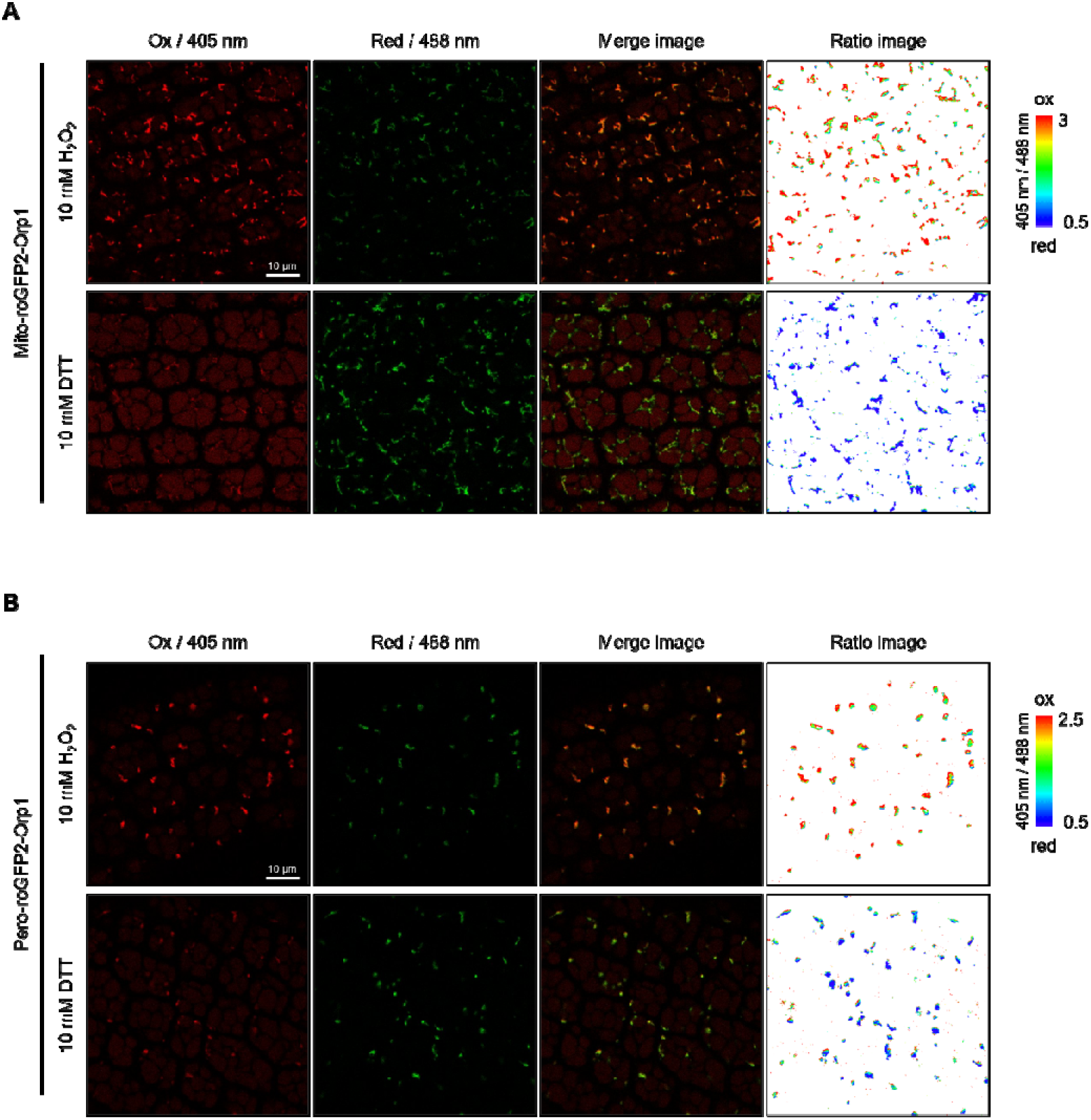
Calibration of the Mito-roGFP2-Orp1 and Pero-roGFP2-Orp1 redox sensors in Arabidopsis seeds. Calibration of Arabidopsis seeds expressing Mito-roGFP2-Orp1 (**A**) or Pero-roGFP2-Orp1 (**B**). Biosensors were fully oxidized by treatment with 10 mM H_2_O_2_ (Ox) or fully reduced with 10 mM DTT (Red). Representative fluorescence images acquired following excitation at 405 nm and 488 nm, merged images, and the corresponding false-color ratio images (405/488) are shown. Scale bars = 10 _μ_*m*.

**Figure S2.**
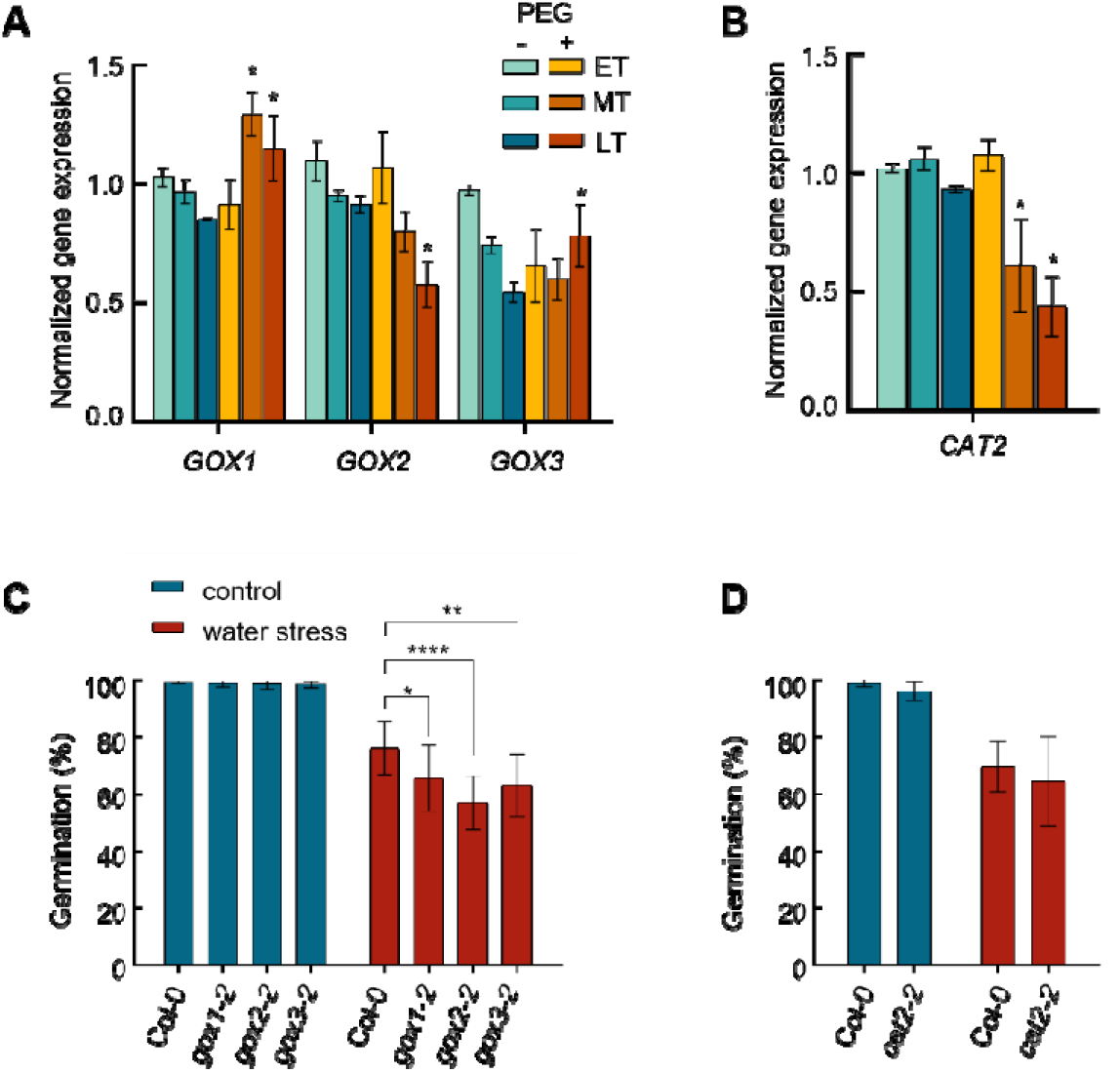
Peroxisome-derived ROS contribute to the germination response to water stress. Transcript abundance of genes involved in peroxisomal ROS metabolism, including ROS-producing enzymes (GOXs, **A**) and ROS-scavenging enzymes (CAT2, **B**), during germination under control and water stress conditions. Data are presented as the mean ± SD of three biological replicates. An asterisk indicates genes that were significantly differentially expressed between control and water stress conditions (FDR < 0.01). Germination assays of loss-of-function mutants affected in peroxisomal ROS production (**C**) or ROS scavenging (**D**) under control and water stress conditions. Data are presented as the mean ± SD of five biological replicates. Statistical significance between treatments at each time point was determined using Student’s t-test. Asterisks indicate significant differences between treatments (P < 0.05; *P < 0.01; ***P < 0.0001).

**Figure S3.**
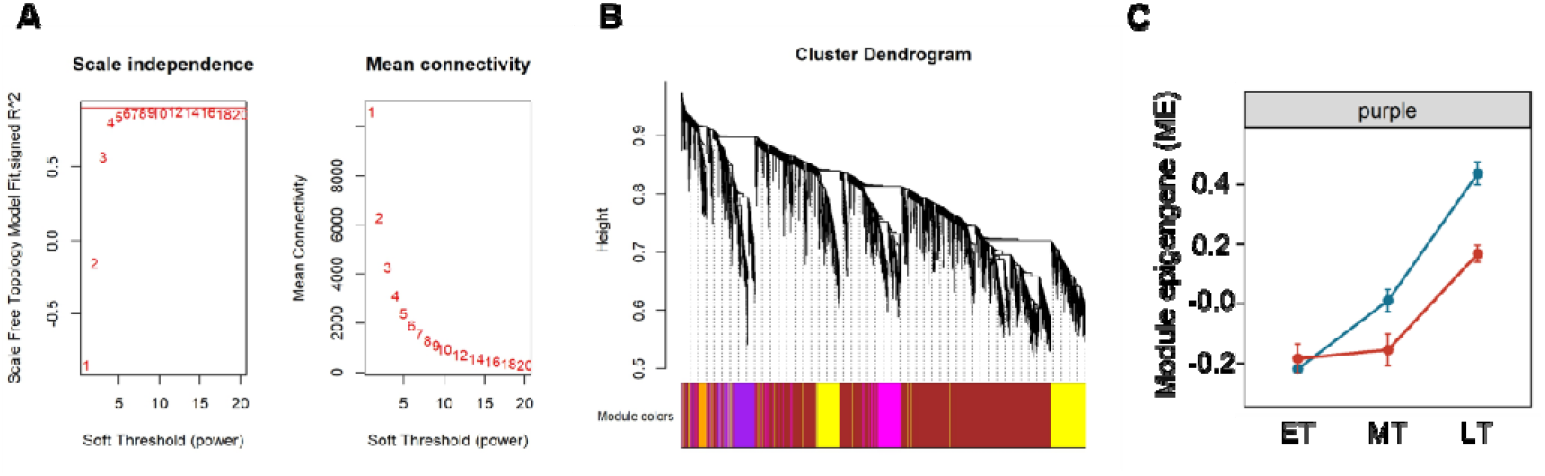
Identification of hub genes by weighted gene co-expression network analysis (WGCNA). **(A)** Analysis of the scale-free topology fit index and mean connectivity as a function of the soft-thresholding power (_β_). **(B)** Hierarchical clustering dendrogram of 24,640 transcripts and the corresponding module assignments. Different colors represent distinct co-expression modules (soft-thresholding power _β_ = 6; scale-free topology fit index, R² > 0.9). **(C)** Expression profile of the purple module eigengene under control (blue) and water stress (red) conditions during germination. Data are presented as the mean ± SD of normalized transcript counts from three biological replicates. ET, early time point; MT, middle time point; LT, late time point.

Table S1: Differentially expressed genes under water stress condition vs control condition at ET

Table S2: Differentially expressed genes under water stress condition vs control condition at MT

Table S3: Differentially expressed genes under water stress condition vs control condition at LT

Table S4: ROS-related transcriptional signatures associated with the response to water stress during seed germination.

Table S5: Gene list of purple module acquired by gene set enrichment analysis

